# Interactions between single actin and vimentin filaments

**DOI:** 10.64898/2026.06.11.731581

**Authors:** Pallavi Kumari, Sascha Lambert, Komal Bhattacharyya, Kristian A. T. Pajanonot, Stefan Klumpp, Sarah Köster

## Abstract

The cytoskeleton determines cell shape, mechanical properties, and motility by interconnected networks of protein filaments – actin filaments, microtubules and intermediate filaments. Their collective function relies on crosstalk between these filament systems, yet the physical basis of interactions between the filaments remains insufficiently understood. Actin and vimentin filaments and networks frequently co-localize within cells and jointly regulate contractility, force transmission and mechanical resilience, indicating functional cooperation. However, it remains unclear whether these interactions arise from direct filament–filament interactions or are mediated exclusively by accessory crosslinking proteins. Studies of reconstituted composite networks probing direct interactions by rheology have yielded inconsistent results. Here, we show that single actin filaments and vimentin intermediate filaments do indeed interact directly, forming force-bearing contacts in the absence of crosslinking proteins, with interaction strengths comparable to other previously reported cytoskeletal filament pairs. Using quadruple optical tweezers combined with microfluidics and confocal microscopy, we systematically probe these interactions under controlled conditions across a range of ionic environments. We find that, in contrast to other filament pairs, variations in ionic strength do not appreciably affect the interaction breaking forces between actin and vimentin filaments. However, the interaction geometry determines the achievable interaction strengths, because the limited stretchability of the actin filaments sets an upper bound to the measurable force range. This limit also imposes a restriction on direct quantification of interaction parameters, which we circumvent by a Bayesian unmasking strategy that allows us to infer bond parameters despite the breaking of actin filaments. Furthermore, actin bundling enhances the stability against breaking, enabling the detection of higher interaction forces. These findings demonstrate that actin and vimentin form a mechanically interacting system through direct filament bonds, and our work establishes a minimal, protein-linker-independent physical basis for actin–vimentin crosstalk and quantifies the interaction forces between the two filament types.

## Introduction

The structure, mechanics and function of eukaryotic cells are largely determined by dynamic cytoskeletal networks comprising actin filaments, microtubules, and intermediate filaments (IFs).^1–3^ Many essential cellular processes, including division and migration, as well as mechanical integrity, critically depend on interactions between these filaments.^4,5^ For instance, actin filaments and microtubules jointly regulate cell shape and polarity.^6^ Microtubules and vimentin IFs interact with kinesin-1 and dynein motors, thereby organizing vimentin precursors and mature filaments along microtubule tracks.^7–9^ While microtubules initially template vimentin assembly, the resulting long-lived vimentin network reciprocally guides microtubule growth as a spatially organized template, ^10^ and directly stabilizes dynamic microtubules by suppressing catastrophes and promoting rescues through hydrophobic and electrostatic interactions.^11^ In specific cell types, keratin IFs organize into a subplasmalemmal rim directly beneath the actin cortex, where their concerted action regulates cellular mechanics and guarantees structural integrity when the tissue is subjected to high strain.^12–14^ The actin cortex also interacts with vimentin IFs, the most abundant IF typically expressed in mesenchymal cells.^15–17^ During mitosis vimentin IFs redistribute to the cell periphery and integrate into the actin cortex, and thereby modify the actin organization in a vimentin tail domain-dependent manner.^18^ Specifically, once recruited by the crosslinking protein plectin, vimentin leads to thinning of the actin cortex and helps generate the high mechanical tension required for the cell to round up for division.^19^ Additionally, actin and vimentin form interpenetrating networks in fibroblasts that significantly increase the cell’s contractility and accelerate recovery after mechanical stress. ^20^

Despite these clear functional synergies between actin and vimentin, it remains unresolved whether the two types of cytoskeletal filaments interact directly or whether their interplay relies entirely on crosslinking proteins. Interestingly, previous rheology studies on reconstituted composite networks of actin and vimentin led to diverging results. Early work by Esue et al. indeed reported direct interactions mediated by the vimentin tail domain, resulting in synergistic stiffening, i.e., the mixed network exhibiting higher stiffness than networks containing either actin or vimentin alone.^21^ In contrast, Golde et al. concluded that these networks behave as a superposition of two non-interacting scaffolds, as they found no evidence of emergent mechanical effects.^22^ More recently, Shen et al. found that vimentin primarily increases the relaxation times and promotes solid-like behavior, while actin determines the overall elastic modulus.^23^ All cytoskeletal filaments are negatively charged polyelectrolytes and their interactions are therefore highly sensitive to the ionic environment.^11,24–29^ The discrepancies between previous rheology studies on mixed actin–vimentin networks may thus arise from differences in the buffer compositions used.

To investigate this hypothesis, we adopt a reductionist approach and study a “minimal network” of only two individual filaments. Using a quadruple optical tweezer setup, we bring an actin and a vimentin filament into contact and move them with respect to each other. ^11,29^ This approach enables us to quantify the interaction forces and binding rates while systematically varying the ionic conditions. We observe that actin and vimentin filaments indeed interact directly even in the absence of crosslinking proteins, and their interaction strength is comparable to previously reported vimentin—vimentin and vimentin—microtubule filament pairs.^11,29^ Furthermore, we employed simulations and Bayesian inference to quantitatively characterize the unbinding rates between actin and vimentin filaments. Since frequent breaking of the actin filament limits the experimental observation of the actin—vimentin interaction under high forces, we develop a Bayesian unmasking strategy that allows us not only to infer interaction parameters from the experimental data despite this limitation, but also to computationally reconstruct experimental results in the absence of actin breaking. Thus our results demonstrate a direct interaction between actin and vimentin filaments and establish a minimal, protein-linker-independent physical basis for actin—vimentin crosstalk.

## Materials and Methods

### Vimentin purification, labeling and assembly

Vimentin purification, labeling, and assembly are performed following established protocols.^30–32^ Briefly, recombinant vimentin C328N, carrying a C-terminal extension of two glycine residues and a cysteine residue to enable site-specific labeling, is expressed in *E. coli*, purified from inclusion bodies, and stored at −80*^◦^*C in denaturing buffer (8 M urea (Bio-science grade; Carl Roth, Karlsruhe, Germany), 1 mM EDTA (ethylenediamine tetraacetic acid; Carl Roth, Karlsruhe, Germany), 0.1 mM EGTA (ethylene glycol-bis(*β*-aminoethyl ether)-N,N,N*^′^*,N*^′^*-tetraacetic acid; Carl Roth, Karlsruhe, Germany), 10 mM methylamine hydrochloride (Sigma-Aldrich, Darmstadt, Germany), 0.15–0.25 M KCl (potassium chloride; Carl Roth, Karlsruhe, Germany), 5 mM Tris (pH 7.5; Carl Roth, Karlsruhe, Germany)). After thawing, vimentin monomers are labeled via maleimide chemistry using ATTO647N (ATTO-TEC GmbH, Siegen, Germany) for fluorescence, and biotin (Sigma-Aldrich, Darmstadt, Germany) for attachment to streptavidin-coated beads. Labeled and unlabeled monomers are mixed to obtain a final composition of 3% fluorescently labeled and 10% biotinylated protein. Renaturation and formation of dimers and tetramers are achieved by stepwise dialysis (using 50 kDa MWCO tubing from Spectra/Por 7, Carl Roth, Karlsruhe, Germany), reducing the urea concentration from 8 M to 0 M in a step-wise manner (8, 6, 4, 2, 0 M) in 2 mM phosphate buffer (pH 7.5; Sigma-Aldrich, Darmstadt, Germany). Filament assembly is induced at a protein concentration of 0.2 mg/mL by adding 100 mM KCl and incubating at 36*^◦^*C overnight to allow for the formation of long filaments. Prior to the experiments, the filaments are diluted 1000-fold in assembly buffer (2 mM phosphate buffer, pH 7.5, supplemented with 100 mM KCl).

### Actin purification, labeling and assembly

Actin preparation is carried out following established protocols.^33–35^ Monomeric actin (G-actin, 42 kDa) purified from rabbit skeletal muscle is stored at −80*^◦^*C in G-buffer (5 mM Tris-HCl (tris(hydroxymethyl)aminomethane hydrochloride; Carl Roth, Karlsruhe, Germany), pH 8.0, 0.2 mM ATP (adenosine 5’-triphosphate; Sigma-Aldrich, Darmstadt, Germany), 0.2 mM CaCl_2_ (calcium chloride; Carl Roth, Karlsruhe, Germany), and 0.5 mM DTT (dithiothreitol; Carl Roth, Karlsruhe, Germany), which maintains actin in its monomeric, ATP-bound state. Biotinylated actin is purchased (Cytoskeleton Inc., Denver, Colorado, USA) and is reconstituted in the same buffer. For fluorescent labeling, G-actin is first dialyzed (25 kDa MWCO tubing; Spectra/Por 7, Carl Roth, Karlsruhe, Germany) overnight at 4*^◦^*C into HEPES (4-(2-hydroxyethyl)-1-piperazineethanesulfonic acid; Carl Roth, Karlsruhe, Germany)-based G-buffer (5 mM HEPES, pH 8.0, 0.2 mM ATP, 0.2 mM CaCl_2_, 0.5 mM DTT) to remove Tris, which interferes with NHS-ester chemistry. Actin is polymerized by adding 1 part 10× F-buffer (100 mM HEPES, 500 mM KCl, 20 mM MgCl_2_ (Carl Roth, Karlsruhe, Germany), 20 mM ATP, pH 7.5) to 9 parts G-buffer and incubating for 1 h at room temperature at 0.03 mg/mL. Fluorescent labeling is performed on F-actin using ATTO488 NHS-ester (ATTO-TEC GmbH, Siegen, Germany) at an 8-fold molar excess relative to actin for 1 h at room temperature, followed by quenching with 10 mM Tris, pH 7.8. Polymerization-incompetent actin is removed by ultracentrifugation at 100,000×g for 1 h at 18*^◦^*C. The pellet is resuspended and depolymerized on ice for 1 h, followed by a second polymerization–ultracentrifugation cycle and dialysis into Tris-based G-buffer. A final ultracentrifugation step (100,000×g, 3.5 h, 4*^◦^*C) removes residual oligomers, and labeled actin is flash-frozen in liquid nitrogen and stored at −80*^◦^*C. For filament assembly, labeled, biotinylated, and unlabeled actin are thawed and mixed to obtain final fractions of 10% fluorescently labeled and 30% biotinylated monomers at a total concentration of 0.2 mg/mL. The mixture is incubated on ice for 2 h to depolymerize pre-existing oligomers. Polymerization is then performed as described above using Tris-based buffers. The resulting F-actin is diluted to a final concentration of 0.003 mg/mL (70 nM) in the assembly buffer and is stabilized with an equimolar concentration of phalloidin (Fisher Scientific GmbH, Schwerte, Germany). For optical tweezer experiments, the filaments are further diluted 100-fold in assembly buffer prior to use.

### Crossed-filaments experiments

Optical tweezer experiments are performed using a commercial system (LUMICKS, Amsterdam, The Netherlands) including quadruple optical traps, a microfluidic flow cell, and confocal microscopy, as described previously. ^11,29^ Briefly, the system is equipped with a 1064 nm (21 W) trapping laser and a 60× water-immersion objective (NA 1.2; Nikon CFI Plan Apo). Fluorescence excitation is performed using 488 nm and 638 nm lasers. The experiments are carried out in a four-channel microfluidic chip, in which each inlet contains a specific component required for filament manipulation and measurement (see Figure 1a). The channels contain: (i) streptavidin-coated polystyrene beads (4.38–4.83 *µ*m; 0.5% w/v; Kisker Biotech GmbH, Steinfurt, Germany) diluted 100-fold in vimentin assembly buffer, (ii) actin filaments in actin assembly buffer, (iii) vimentin IFs in vimentin assembly buffer, and (iv) a common measuring buffer (CB) containing 100 mM KCl, 1 mM MgCl_2_, and 2 mM phosphate buffer (pH 7.5), supplemented with an oxygen scavenging system (1.2 mg/mL glucose, 0.02 mg/mL glucose oxidase, 0.004 mg/mL catalase, 20 mM DTT) and 0.01 mM phalloidin. The ionic conditions of the measuring buffer are systematically varied. This includes increasing the MgCl_2_ concentration to 5 mM or 20 mM, reducing the KCl concentration to 90 mM (without divalent ions), and adding 0.1% (w/v) Triton X-100 (TX100; Carl Roth, Karlsruhe, Germany). An intermediate channel containing vimentin assembly buffer is placed between the measuring buffer and vimentin channels for conditions with ionic strength above CB, enabling precise capture of single filaments and attachment to the bead pair without bundling. For additional divalent cation experiments, the phosphate buffer is replaced by 10 mM HEPES (pH 7.5) to avoid cation–phosphate interactions. 19 mM MgCl_2_, CaCl_2_, or MnCl_2_ (Merck Chemicals GmbH, Darmstadt, Germany) are added to HEPES-CB with 0.5 mM EGTA. DTT is replaced by Tris(2-carboxyethyl)phosphin (TCEP) to avoid interference with divalent ions. An intermediate channel containing actin assembly buffer without oxygen scavenger or reducing agent, placed between the channels for the measuring buffer and the vimentin filaments, prevents buffer cross-contamination and unwanted reactions between phosphate buffer and divalent ions at high concentrations, especially of MnCl_2_. All solutions are filtered through 0.2 *µ*m pore size cellulose acetate membranes (Th. Geyer, Renningen, Germany) before use. For each experiment, four beads are trapped and arranged into two bead pairs (Figure 1bI). One pair is used to capture a single vimentin IF, and the other one to capture a single actin filament (Figure 1bII,III). After transfer to the measuring buffer, the filaments are arranged in a crossed geometry and are adjusted such that both filaments lie in the same focal plane (Figure 1bIV). The interaction measurements are performed by displacing one filament with respect to the other one at a speed of 0.27 ± 0.03 *µ*m/s. A pause of 1 s is applied between displacements. The forces on bead 1 (Figure 1c) are recorded in the *x*- and *y*-directions, while confocal images are acquired simultaneously. The measurements are continued until a filament—filament interaction or a filament itself breaks.

**Figure 1:**
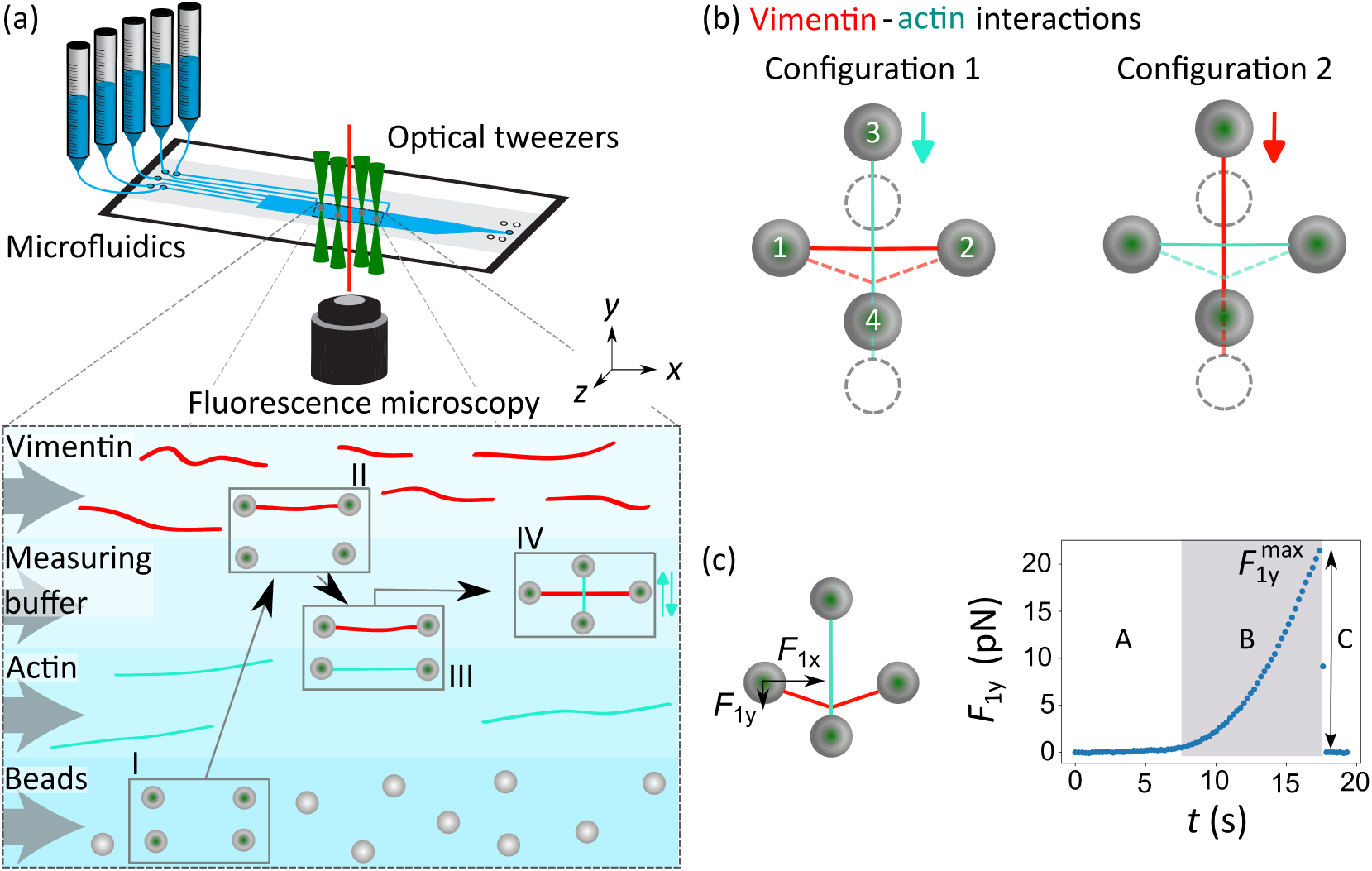
(a) (Top) Overview of the experimental setup combining quadruple optical tweezers, a microfluidic chip for laminar co-flow, and confocal fluorescence microscopy. (Bottom) Schematic of the experimental workflow within the microfluidic chip. (b) Two different measurement geometries: (Left) Configuration 1 with the actin filament pulled across the vimentin filament, which consequently is extended; (Right) Configuration 2 with the vimentin pulled across the actin filament, which consequently is extended. (c) (Left) Schematic force diagram. The *x* and *y* components of the force acting on bead 1 is detected. (Right) The *y* component *F*_1*y*_ plotted versus the time *t*; in regime A the force is zero, it increases in regime B and suddenly drops to zero in regime C. The force *F* ^max^ provides the maximum interaction force between the two filaments.

### Single actin filament pulling experiments

Optical tweezer experiments are performed in a microfluidic flow cell as previously described in Refs.,^31,36^ using a three-channel configuration. The channels contain (i)streptavidin-coated beads in actin assembly buffer, (ii) single actin filaments in actin assembly buffer, and (iii) measuring buffer. All experiments are performed under the same measuring buffer conditions, as described for the crossed-filaments experiments, to ensure comparability across measurements. Single actin filaments are first captured between two optically trapped beads (beads 1 and 2) by transferring the bead pair into the actin channel until a single filament binds between the beads. After successful attachment, the bead–filament ensemble is moved into the measuring buffer channel for force-extension measurements. For the filament stretching experiments, trap 1 is kept fixed while trap 2 is displaced along the filament axis (positive *x*-direction), thereby applying a longitudinal force to the actin filament (see Figure 3b). The displacement is performed at a speed of 0.27 ± 0.03 *µ*m/s. Force and extension are recorded continuously during stretching. Each measurement is continued until the filament breaks.

### Data analysis

Confocal image analysis and all force data processing are performed using custom-written Jupyter Notebook scripts. For the crossed-filaments experiment, the analysis follows Refs. ^11,29^ Briefly, the *y*-component of the force acting on bead 1 (*F*_1*y*_) is analyzed, as forces in the *x*-direction are balanced (see Figure S1). Interaction events are manually identified from the raw force–time data, followed by background subtraction. Since trap 1 (and thus bead 1, defined as the bead held in trap 1) is not actively being moved during the experiment, the forces are determined from the displacement of bead 1 relative to the trap center. To calculate the total breaking force, the measured force *F*_1*y*_ is multiplied by a geometric correction factor (see Figure S1), which depends on the distance between the horizontal beads (center to center) and the distance between the center of the left bead and the contact point of the crossed filaments. The mean binding rate is estimated by bootstrap resampling following the approach described in Ref.,^29^ because the binding events are rare.

For the single filament stretching experiments, the analysis is conducted similarly to Refs.^29,36^ The recorded *F*_1*x*_ force–extension trace is background-corrected. The strain, *ε*, is calculated from the filament length:

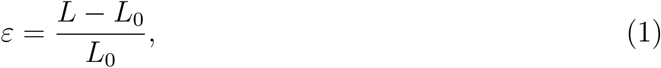

where *L* is the instantaneous filament length and *L*_0_ is the initial filament length measured at a force of 1 pN.

### Two-bond model

We describe the actin–vimentin system by a minimal two-bond model consisting of (i) the actin—vimentin interaction (index ‘int’) and (ii) a solitary weak link within the actin filament (‘act’). Both bonds are modeled as independent two-state systems (bound/unbound) with force-dependent Bell-Evans unbinding kinetics, ^37,38^

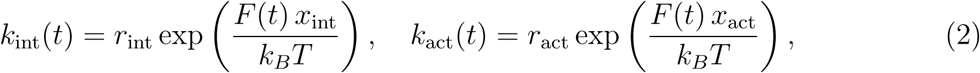

with two parameters for each bond, the force-free unbinding rate *r*_int,act_ and the distance *x*_int,act_ to the transition state that characterizes the force sensitivity of the bond. Under the experimental protocol in configuration 1, both bonds experience the same time-dependent force *F* (*t*) and can be thought of as being arranged in series. Rupture of the two bonds thus constitutes two competing processes and the system evolves under a time-dependent loading protocol until the first rupture event occurs. The survival probability up to time *t* is 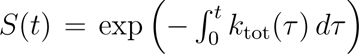 with the total rupture rate *k*_tot_(*t*) = *k*_int_(*t*) + *k*_act_(*t*) and the probability density for rupture at time *t* is *p*(*t*) = *k*_tot_(*t*) *S*(*t*). At that time, the actin— vimentin interaction breaks with probability *k*_int_(*t*)*/k*_tot_(*t*) and the actin filament with 1 − *k*_int_(*t*)*/k*_tot_(*t*).

Conceptual simulations of the model are performed with a time-dependent force *F* (*t*) following an explicit polymer model for stretching of the filament.^11^ In contrast, to obtain quantitative predictions to be compared with the experiments and counterfactual simulations without breaking of actin, we use predictive posterior simulations based on the Bayesian inference described below. These simulations (as well as Bayesian inference itself) use the experimental force trajectories *F* (*t*) directly, and thus do not depend on a polymer model for filament stretching. For details of both types of simulations, see the Supplementary Material.

### Bayesian inference of bond parameters

To estimate the parameters of the two-bond model *θ* = (*r*_int_, *x*_int_, *r*_act_, *x*_act_) from the experimental observations, including the trajectories terminated by actin rupture, we use a Bayesian inference framework. The inference jointly estimates the kinetic parameters of the actin—vimentin interaction and the effective actin weak-link description from the measured force-time trajectories *F* (*t*). Each data point (*t_i_*, *F_i_*) is assigned a discrete rupture label *y_i_* (= act, int, none) to indicate which bonds break. Generalizing the approach from Ref.^39^ to two competing rupture processes, we obtain an explicit expression for the likelihood L(data|*θ*) as a product over independent observations, also including the data of single actin pulling experiments with the corresponding one-bond model. Combined with uniform priors *P* (*θ*) for the parameters, this provides the posterior distribution *P* (*θ*|data) ∝ L(data|*θ*) *P* (*θ*) of the four parameters, which is sampled using an affine-invariant ensemble Markov Chain Monte Carlo method.^40^ Details of the likelihood, priors, and sampling procedure are provided in the Supplementary Material.

## Results

### Actin and vimentin filaments interact directly

The quantification of inter-filament interactions requires precise force detection as well as visual control over the experiment and a versatile sample environment. Furthermore, to avoid substrate interactions, measurements in solution are desirable. To meet these requirements, we employ an integrated experimental setup that combines quadruple optical tweezers for force measurements, confocal fluorescence microscopy for simultaneous imaging, and a microfluidic sample environment for tuning the interaction conditions.^11,29^ A schematic of the setup is shown in Figure 1a, top. Below, a schematic top view of the microfluidic flow cell is shown. It includes four separate pressure-driven inlets that generate laminar co-flow of four solutions within the channel, used for vimentin filaments, measuring buffer, actin filaments and beads. The experimental workflow is indicated by the Roman-numbered rectangles. (I) Four streptavidin-coated beads are captured by the four optical traps. The stiffness of each trap is calibrated by analyzing the power spectral density of the thermal fluctuations of the bead in the measuring buffer channel prior to filament attachments. (II) Fluorescently and biotin-labelled vimentin filaments are first attached to one bead pair via biotin–streptavidin binding, (III) followed by attachment of actin filaments to a second bead pair. (IV) Both filaments are transferred to the measuring buffer channel and arranged in a crossed geometry within the same *z*-plane for interaction measurements. For simplicity, we refer to the two filaments as “horizontal” and “vertical” according to their orientation in the image, corresponding to the *x*- and *y*-directions, respectively.

As we investigate interactions between two different types of filaments, two principal geometric configurations are possible with (1) the actin filament positioned vertically and the vimentin filament positioned horizontally, and (2) vice versa, see Figure 1b, left and right, respectively. To probe filament–filament interactions, the vertical filament is repeatedly moved upwards and downwards perpendicularly to the horizontal filament until an interaction forms and eventually ruptures. Once an interaction between the filaments forms, the displacement of the vertical filament with respect to the horizontal filament causes the horizontal filament to bend and extend (see dashed lines in Figure 1b). With further displacement, the bond ruptures and both filaments relax back to a straight appearance. During the vertical movement of traps 3 and 4, the forces acting on bead 1 (Figure 1c, left) are measured. An example force–time plot (Figure 1c, right) exhibits three regimes: (A) zero force when the filaments are unbound, (B, shaded) a gradual increase during bond formation as the vertical filament is displaced, (C) and a sudden drop indicating the bond rupture. The interaction strength is the maximum force directly before the force drops to zero, termed “interaction breaking force” (*F* ^int^). This force is calculated from the maximum measured force on bead 1 (*F* ^max^) and the filament geometry at the interaction site (see Figure S1).^11,29^

Our setup allows for confocal imaging simultaneously with the quadruple optical tweezers experiments. We are thus able to directly visualize the interactions between single actin and vimentin filaments. Figure 2a,i shows example time-lapse snapshots of a phalloidin-stabilized actin filament (cyan, vertical) crossing a single vimentin filament (red, horizontal) (configuration 1). As the actin filament is moved downward (cyan arrow), the horizontal vimentin filament bends, providing a clear qualitative indication that the filaments directly interact. The actin filament exhibits an asymmetric response. The lower segment (highlighted by the yellow rectangle) becomes brighter as it experiences tension and moves in focus, whereas the upper segment (highlighted by the magenta rectangle) relaxes and slightly moves out of focus. Due to the low tolerance of actin filaments to tension, they need to be kept slightly relaxed prior to the interaction measurements. Therefore, and in contrast to vimentin filaments that can sustain substantially higher tensile loads without breaking, ^31^ we cannot pre-stretch the actin filaments to pull out the entropic fluctuations.^11^ Upon further vertical displacement of the actin filament, the actin–vimentin interaction breaks, indicated by the cyan double arrow. In other cases, the actin filament itself breaks (cyan star) while the actin–vimentin bond remains intact (Figure 2a,ii). Swapping the orientations of the filaments (configuration 2) results in the same two outcomes, i.e., interaction breaking and filament-breaking, as shown in Figure 2a,iii. If a filament breaks, it is typically the actin filament and only in rare cases the vimentin filament.

**Figure 2:**
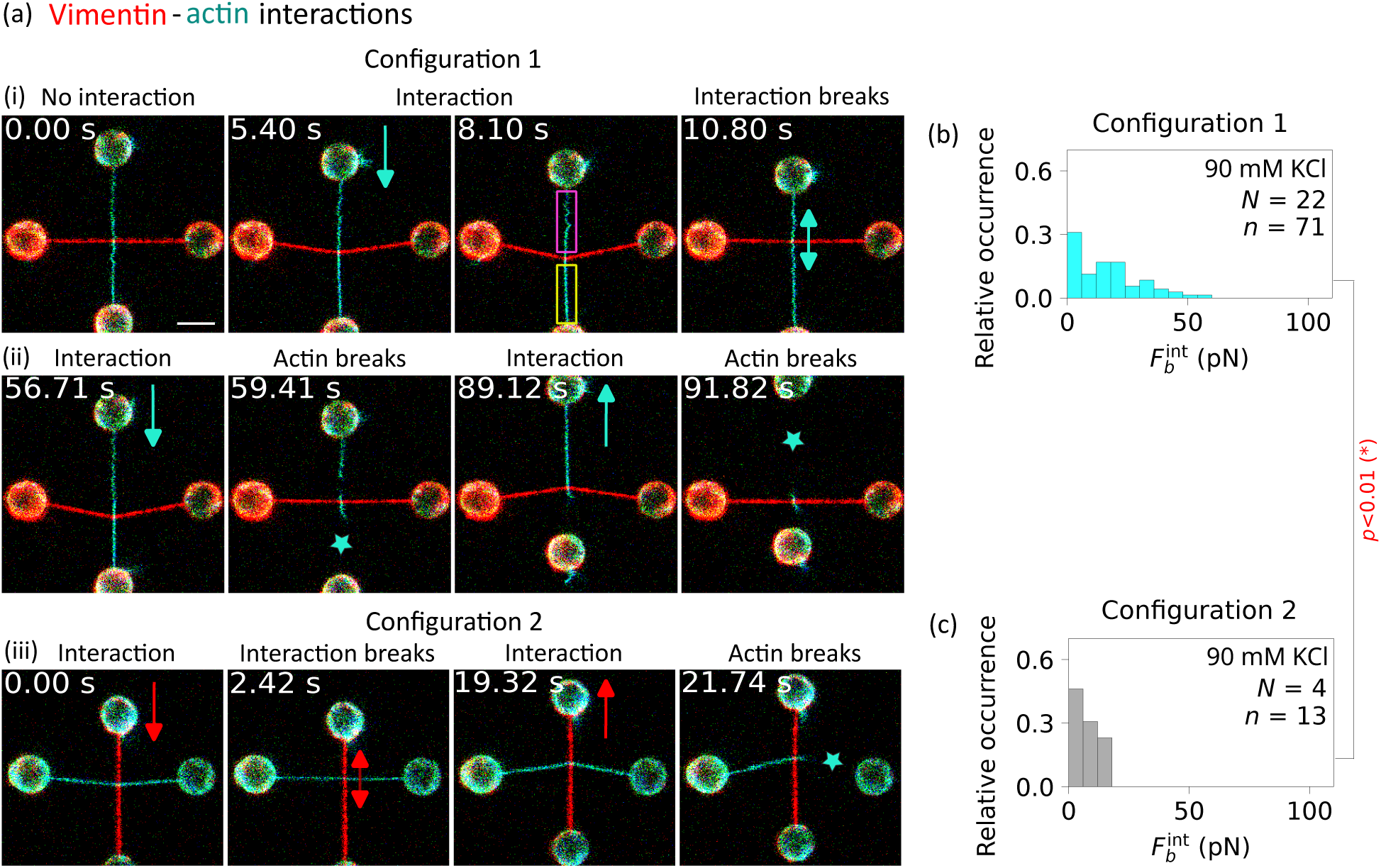
(a) Fluorescence images of typical interaction events; shown are snapshots from experimental videos (see Supplemental Movies S1, S2, and S3); (i) the vimentin filament (horizontal) gets extended and the filament interaction breaks (cyan double arrow); (ii) the vimentin filament (horizontal) gets extended and the actin filament breaks (cyan star); (iii) the actin filament (horizontal) gets extended and first the filament interaction breaks (red double arrow) and thereafter the actin filament breaks. Scale bar: 5 *µ*m. (b) Histogram of interaction breaking forces from configuration 1 and (c) from configuration 2. The number *N* of interacting filament pairs and the number *n* of interaction events is shown in the panels. To compare the distributions, the *p*-value is calculated using a two-sample KS test with a significance level of 5% followed by a Holm–Bonferroni correction; red indicates a significant difference.

Figure 2b,c show the interaction breaking force distributions for both configuration 1 and 2, respectively. Here, we study an equal number of interacting filament pairs for each configuration. A single filament pair can undergo multiple interaction—rupture cycles, resulting in a total number of interaction breaking events *n* that exceeds the number of interacting filament pairs *N*. In cases where the actin filament ruptures before the filament–filament interaction, the number of interaction breaking events is limited, resulting in a lower total number of recorded events *n*. The numbers of interaction events and the corresponding force distributions show clear differences between the two configurations. In configuration 1, a substantially higher number of interaction breaking events is observed, and the breaking forces range up to approximately 50 pN, clearly exceeding both the number of events and the force values measured in configuration 2. A two-sample Kolmogorov–Smirnov (KS) test confirms that the force distributions differ significantly between the two configurations. We assign this observation to the higher propensity of actin filaments to break in configuration 2.

Together, these observations highlight the importance of the structural integrity of the filaments in interpreting interaction measurements and in selecting experimental geometries that minimize filament damage, thereby allowing for a wider range of interaction breaking forces to be captured. Accordingly, we focus on configuration 1 to conduct further studies to systematically examine the effects of ion concentration, ion type, buffer type, and the presence of ionic detergent on the interaction breaking forces. Divalent cations are introduced to examine how electrostatic effects and specific ion-mediated interactions modulate the interaction strength. First, we vary the magnesium (Mg^2+^) concentration between 1 and 20 mM to assess concentration-dependent effects. To further probe ion specificity, Mg^2+^ is substituted with other divalent ions, i.e., calcium (Ca^2+^) or manganese (Mn^2+^) while maintaining a total divalent ion concentration of 20 mM. In addition, experiments are performed in different buffer types, i.e., HEPES and phosphate buffer, to test potential buffer-dependent effects. To test whether hydrophobic interactions contribute to filament binding, an ionic detergent (0.1% w/v TX100) is added. Across all these conditions, the measured force distributions remain largely unchanged (see Figure S2), indicating that with our current measurement geometry, we cannot detect any influence of the sample environment on the interaction strength between actin and vimentin filaments.

We further investigate the binding rate between these filament pairs (see Figure S3). The binding rate, *r*_b_, is calculated by dividing the number of interaction events by the total time during which the two filaments remain unbound. For the buffer conditions shown in Figure 2, *r*_b_ is approximately 0.02 s*^−^*^1^, which is comparable to values reported for other filament pairs at similar ionic strength. ^11,29^ Upon increasing the ionic strength through magnesium addition and higher magnesium concentrations, as well as with calcium, *r*_b_ remains unchanged (except for a higher value for 5 mM magnesium). In contrast, *r*_b_ increases to 0.07 s*^−^*^1^ in the presence of manganese and 0.05 s*^−^*^1^ in the case of TX100. In fact, in the presence of manganese and TX100, we frequently observe that, following rupture of the actin filament in the crossed-filament configuration, the loose actin filament segment immediately reattaches to the vimentin IF and subsequently overlaps with it along its length. These observations are consistent with enhanced actin—vimentin association and filament bundling, providing independent support for the increased binding rates measured under these conditions.

### Actin filament extensibility limits the interaction forces

As mentioned above, during the measurements of actin—vimentin interactions, we find a higher propensity of the actin filament to break when using configuration 2. To quantify this observation, we compare the relative occurrences of events, i.e., no interaction at all, breaking of the interaction, or breaking of the actin filament, see Figure 3a. Here, all numbers are normalized to the total number of movements of the vertically oriented filament with respect to the horizontally oriented filament in either direction. Among these interaction events, we observe breaking of actin filaments in about half the cases in configuration 1, as compared to more than 80% in configuration 2. As seen in Figure 2a, upon interaction, the horizontal filament bends in the direction of the movement and, therefore, is stretched. Thus, in configuration 2, the actin filament is affected by this extension.

**Figure 3:**
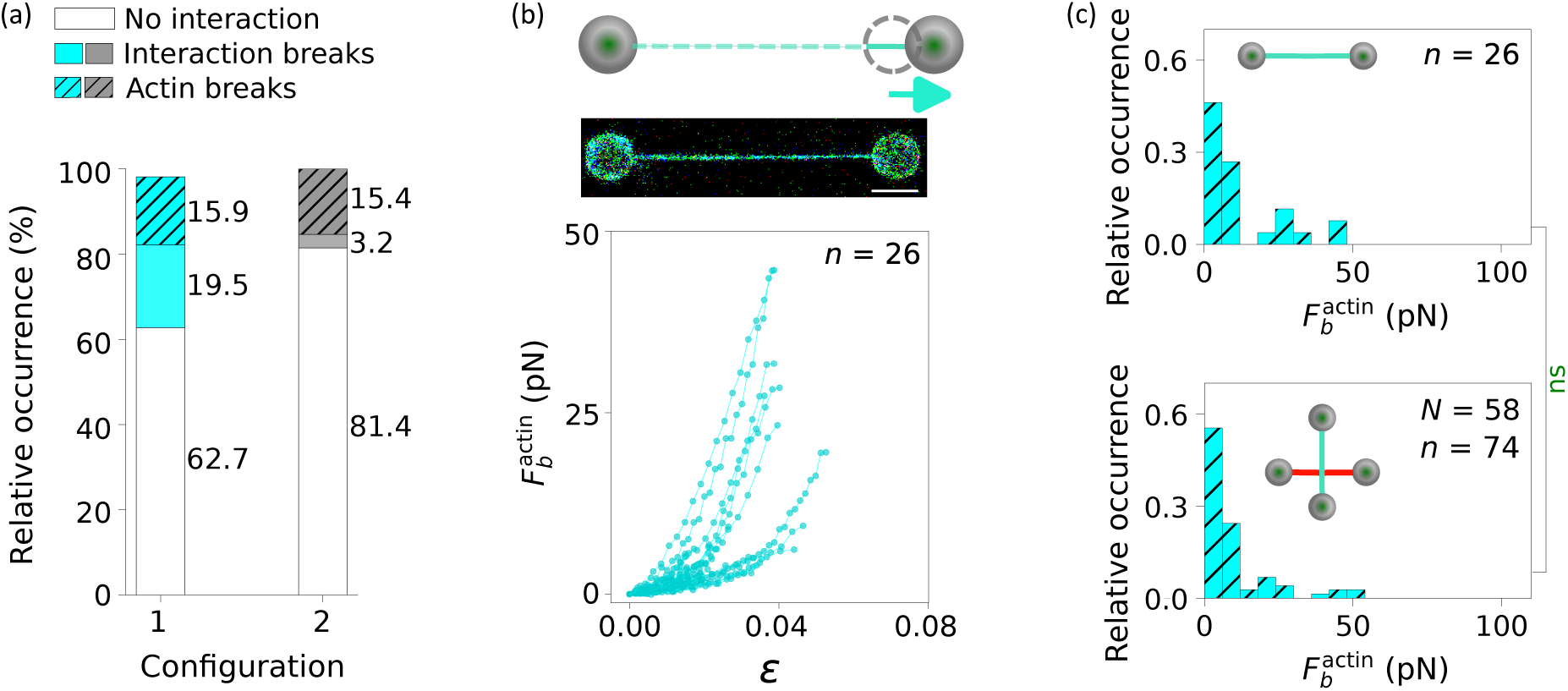
(a) Comparison of the different events occurring for configuration 1 and 2. The data are normalized to total number of movements (up or down) of the vertical filament with respect to the horizontal filament. Note that rare vimentin breaking events have not been included and therefore the values for configuration 1 do not sum up to 100%. (b) Experimental setup including dual optical tweezers for characterizing the force at which an actin filament breaks (top: schematic; center: fluorescence micrograph) and (bottom) force–strain data for individual, stretched actin filaments. Scale bar: 5 *µ*m. (c) Comparison of the actin breaking forces for (top) individual stretched actin filaments and (bottom) breaking events during interaction experiments. The number *N* of filament pairs and the number *n* of actin breaking events is shown in the panels. To compare the distributions, the *p*-values is calculated using a two-sample KS test with a significance level of 5% followed by a Holm–Bonferroni correction; green indicates no significant (ns) difference.

To better understand our observation, we complement the interaction measurements using quadruple optical tweezers by measurements of force-strain data using dual optical tweezers,^31,32,41^ see Figure 3b. Here, single actin filaments are attached to two beads at their ends and stretched until they break, while recording the force *F* and the length *L* to derive the strain *ε*. The recorded force-strain curves reveal that some actin filaments sustain forces around 50 pN, while most of them break at low forces below 20 pN and strains of about 0.03. The result is similar across different buffer conditions (see Figure S4). These low values indicate that actin filaments tolerate only very limited extension under stress, in agreement with Ref.^36^ In contrast, vimentin IFs can withstand forces up to 8 nN and strains up to 3.5, ^31^ and possibly more, highlighting their far greater mechanical resilience. This remarkable difference demonstrates that actin filaments are the mechanically weaker component during filament—filament interactions and are therefore most likely to break. These results also explain why configuration 1 is more suitable for our interaction measurements, as here, primarily the vimentin filament is stretched. When comparing the distributions of actin breaking forces measured by dual optical tweezers filament stretching or by the quadruple optical tweezers interaction assay (see Figure 3c top and bottom, respectively), we find strong agreement, with no significant difference detected between the distributions. This agreement indicates that the filament-breaking events observed in the interaction assay reflect the intrinsic mechanical limits of actin filaments. Experiments using varying buffer conditions (see Figure S5) confirm the robustness of these observations.

It is highly likely that actin filaments are structurally heterogeneous and contain local “predetermined breaking points”. This assumption is supported by cases where, in the crossed-filaments configuration, the actin filament breaks and then reattaches to the bead. Upon further displacement, the actin–vimentin interaction eventually breaks. These data are shown in Figure S6. Quantitative analysis of the forces involved shows that in these cases the interaction breaking forces are always higher than the initial actin breaking forces. We interpret this finding such that once the weak predetermined breaking point is eliminated from the filament, it can resist higher stretching forces.

### Actin bundling extends the range of interaction forces

Since the forces required to break actin—vimentin interactions can exceed the strength of a single actin filament, we next ask whether strengthening the actin filament would influence these interactions. To answer this question, we bundle two actin filaments and investigated their interactions with single vimentin filaments. We therefore perform measurements in buffer containing 19 mM Mn^2+^, which promotes actin bundling by counterion condensation.^24^ Two actin filaments are captured between a bead pair and transferred to the measuring buffer channel (Figure 4a, top). In some cases, bundling occurs spontaneously when the filaments come into contact (Figure 4a, bottom), which is confirmed by confocal fluorescence images. If bundling does not occur spontaneously, the bead pair is gently moved upwards and downwards within the same channel to bring the filaments closer and facilitate bundle formation.

**Figure 4:**
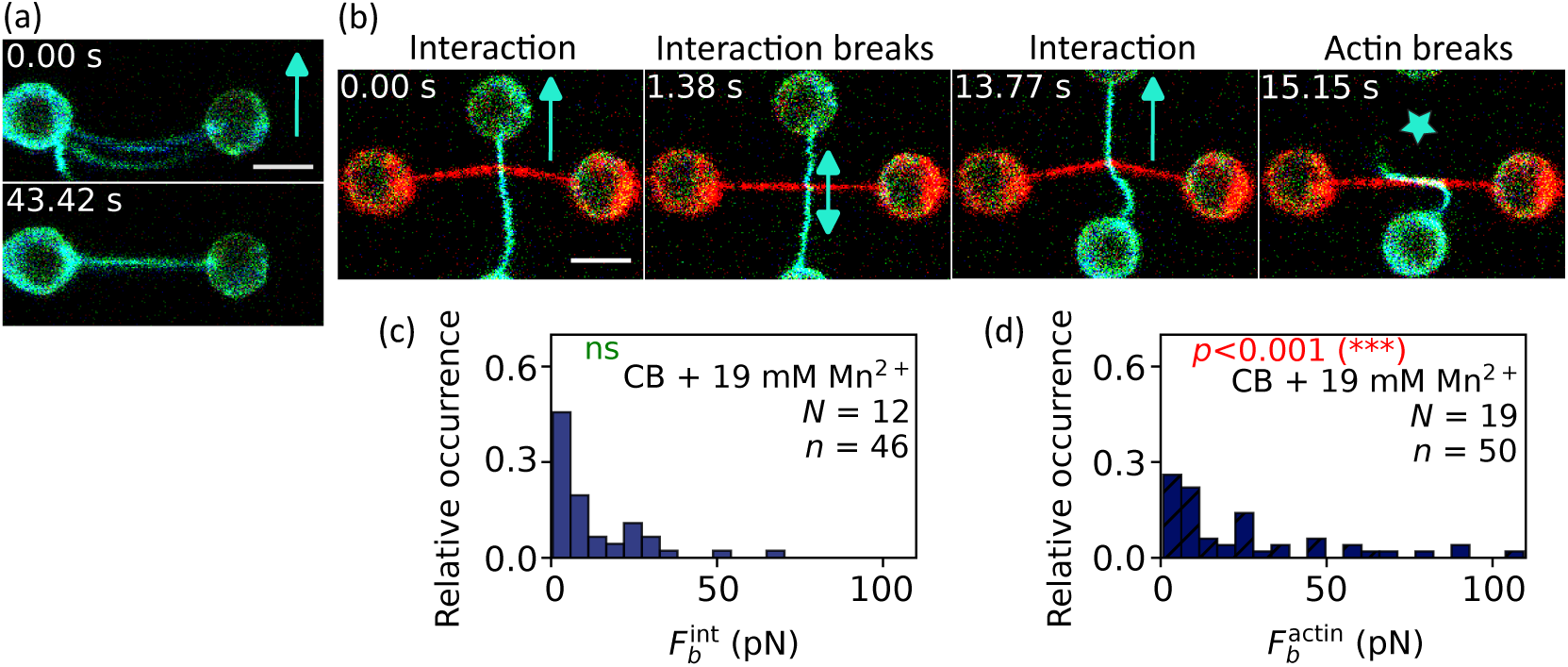
(a) Confocal fluorescence images of two actin filaments (top) that form a bundle (bottom) in the presence of 19 mM Mn^2+^. Scale bar: 5 *µ*m (also for panel b). (b) Confocal fluorescence images showing an interaction between a single vimentin filament and a bundle of two actin filaments; the interaction breaks first, followed by breaking of the actin bundle after several vertical displacements of the vimentin filament. (c) Histogram of interaction breaking forces between a single vimentin filament and a bundle of two actin filaments. (d) Histogram of actin breaking forces of bundles of two actin filaments. The number *N* of filament pairs and the number *n* of actin breaking events is shown in the panels. To compare the distributions, *p*-values are calculated using a two-sample KS test with a significance level of 5% followed by a Holm–Bonferroni correction. The statistical significance compared to the CB is shown inside the panels; green indicates no significant (ns) difference and red indicates a significant difference.

As before for the interactions between single actin filaments and vimentin filaments, we observe two different types of outcomes, i.e., either breaking of the interactions between the actin bundle and the vimentin filaments or breaking of the actin bundle itself (Figure 4b). When comparing the distributions of the interaction breaking forces (Figure 4c) with the case of single actin filaments (Figure 2b), we do not observe a significant difference as shown by the KS test. Thus, interestingly, the bundling and the presence of two actin filaments do not alter the actin–vimentin interaction breaking forces. In contrast, the actin breaking forces change markedly, as shown in Figure 4d. Some of the actin bundles are able to withstand higher forces than single actin filaments, as reflected in a shift of the breaking-force distribution to larger values.

### Bayesian inference unmasks the unbinding force distribution

The quantitative analysis of the breaking of the actin—vimentin bond is not straightforward because of the frequent breaking of the actin filament, which masks the breaking of the actin–vimentin bond. This observation raises the question of whether one can learn parameters of the actin–vimentin bond from the data or whether the experiments predominantly reflect the breaking of actin. To address this question, we describe the actin–vimentin system with a model consisting of two bonds, the actin–vimentin interaction and a weak link in the actin filament, both breaking with Bell–Evans kinetics (see Methods). In configuration 1, the two bonds are subject to the same time-dependent force. They break independently of each other but are, in our experiments, only observed until one of them is broken. In simulations, however, we can continue to observe one bond after the other is broken to illustrate the masking effect on the distribution of the breaking forces. While in some simulations, the actin–vimentin interaction ruptures first, in others, the interaction ruptures only after the breaking of the actin filament, which is not observable experimentally. The full histogram of all interaction rupture events, before and after breaking of actin, gives the characteristic distribution of breaking forces of the interaction that would be observed if actin was stable (light gray bars in Figure S7a,b). Since actin is not stable, only a subset of these rupture events, i.e., those where the interactions break first, are actually observed in the experiments (cyan bars in Figure S7a,b) – the rest of the distribution is masked by actin breaking. Importantly, masking changes the shape of the distribution and suppresses specifically high breaking forces. The extent of masking depends on the parameters of the two bonds, it is particularly pronounced if the actin breaking rate is higher or comparable to the breaking rate of the interaction and if actin is more force sensitive than the interaction (Figure S7c).

To extract parameters of the vimentin–actin interaction from the experimental data, despite the expected masking by the breaking of actin filaments, we turn to Bayesian inference, as our previous work has shown that Bayesian analysis of the full force-time curves can extract more information from this type of experiment than the analysis of breaking force histograms.^39^ Since Bayesian inference provides probability distributions of the parameters rather than just estimates of their values, this approach provides a quantification of the uncertainty and thus indicates how much information our experiments provide about the parameters of the actin–vimentin interaction. Therefore we extended the Bayesian approach from Pajanonot et al. ^39^ to the case of two bonds that can break independently and obtained joint posterior distributions for the four parameters (*r*_int_, *x*_int_, *r*_act_, *x*_act_) of the two bonds based on the the force–time curves from quadruple optical tweezer experiments on crossed actin and vimentin filaments (including trajectories where actin breaks and where the interaction breaks) and the dual optical tweezer experiments with actin only, see Methods and Supplementary Material. Figure 5a,b show two-dimensional marginal distributions of the posterior distribution for the interaction and actin parameters separately (different representations of the full posterior distribution are shown in Figure S8). Figure S9 shows the corresponding plots for the different buffer conditions. The posterior distribution has a similar width in the direction of the actin parameters as in the directions of the interaction parameters, indicating that all four parameters can indeed be inferred. Across the conditions, the force-free unbinding rate is indeed slightly higher for the breaking of the actin filament than for rupture of the actin–vimentin bond. At the same time, the actin breaking is characterized by a slightly lower force sensitivity than the actin–vimentin interaction (*x*_act_ *< x*_int_). The posteriors for different buffer conditions show considerable overlap (Figure S9a,b), confirming the observation that the effects of ions and buffers are small.

**Figure 5:**
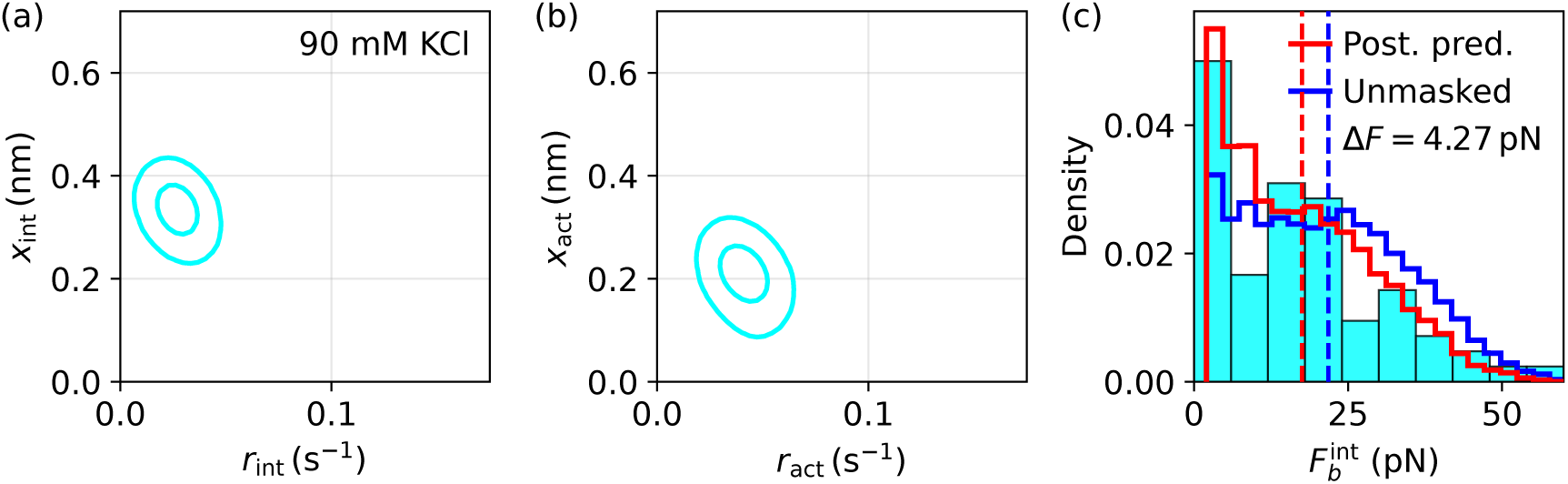
Results of the Bayesian inference on an experimental dataset. (a) and (b) Two slices of the 4D posterior distribution that describe all four kinetic bond parameters. Drawn are the 95% and 50% highest density regions. (c) Empirical distribution of observed breaking forces as a histogram, along with the posterior predictive distribution of this observable from the inferred model (in red) and the counterfactual model with stabilized actin (blue). The distributions show only the portion of results in which the actin does not break (which is not relevant to the stabilized actin prediction). The stabilized actin model shows a shift to higher breaking forces because actin breaking no longer masks long-lived interaction bonds. Vertical dashed lines indicate the prediction center of masses, separated by Δ*F*.

Finally, we used the posterior distribution of the parameters to run predictive simulations to determine breaking force distributions of the actin–vimentin bond, first for the experimental scenario with breaking of actin filaments and then for the desired, but experimentally inaccessible scenario without actin breaking. The posterior prediction of the unbinding force histogram for the experimental scenario (red curve in Figure 5c) agrees well with the experimental results (cyan bars). The distribution in the hypothetical scenario where actin does not break (blue curve) is shifted towards higher forces compared to the case with actin breaking, as also indicated by the vertical dashed lines. This shift is seen consistently across all conditions we studied, as shown in Figure S9c. This comparison confirms that the breaking force distribution of the interaction is indeed masked by actin filament-breaking, but also that the posterior prediction allows us to computationally remove the experimental limitation due to actin breaking and to unmask the breaking force distribution.

## Discussion

The mechanical interplay between cytoskeletal filaments is central to understanding how cells regulate structure and force transmission. In this context, our results place the interaction breaking forces between actin and vimentin filaments, ranging up to ∼50 pN, and therefore within a regime comparable to other cytoskeletal filament pairs, i.e., vimentin–microtubule and vimentin–vimentin filaments.^11,29^ Additionally, these forces are similar to the breaking forces between actin and the specific crosslinkers filamin and *α*-actinin (40 – 80 pN).^42^ This correspondence suggests that the interactions between actin and vimentin filaments are mechanically significant and may contribute to cytoskeletal integrity even without crosslinker proteins. The crosstalk between actin and vimentin filaments likely arises from direct physical interactions, potentially mediated by the vimentin tail domain, in the absence of dedicated crosslinking proteins.^18,20,21^ Furthermore, recent studies reported that vimentin promotes actin assembly by stabilizing ATP-actin at the barbed end, independent of its head and tail domains, indicating multiple interaction interfaces.^43^ The direct interactions provide a molecular basis for the interpenetrating networks observed in the cell cortex and stress fibers.^20^ Rather than functioning as independent systems, actin and vimentin filaments may form a mechanically integrated composite through transient yet sufficiently strong filament-–filament contacts. Consistent with this view, these findings establish a direct link between single-filament interactions and cooperative behavior observed in composite cytoskeletal networks, such as the reported increase in contractile magnitude in fibroblasts containing interpenetrating networks.^20^

Notably, we observed interactions between actin and vimentin filaments and subsequent breaking events in ionic environments containing only monovalent ions as well as in the presence of divalent ions. These findings indicate that direct actin–vimentin interactions are mechanically robust and can form without divalent counterions, suggesting that monovalent electrostatic screening is sufficient to mediate crosstalk between the two filament types. The buffer used for the measurements contains potassium, magnesium, calcium, or manganese ions, which are representative of intracellular environments.^44,45^ Within the studied ionic range, the forces required to break the interactions remain unchanged across different ionic conditions. This distinguishes the interactions between actin and vimentin filaments from those between vimentin–microtubules or between vimentin–vimentin filaments, where divalent ions and changes in ionic strength play a more pronounced role. There is no evidence of significant hydrophobic contributions under the studied conditions, as assessed with TX100 detergent. The direct measurements help clarify the long-standing inconsistencies in actin–vimentin network studies, where some studies describe these systems as weakly interacting^21^ or independent,^22^ while another study reports synergistic mechanical effects, including increased stiffness and altered relaxation dynamics.^23^

On the data analysis side, we have developed a Bayesian unmasking strategy, extending our recently introduced Bayesian analysis framework.^39^ This strategy, which may also be useful in other applications, allows us to infer bond parameters from the experimental data despite the effect of breaking actin filaments, which prevents the observation of most interaction breaking events at high forces. We used posterior predictions to reconstruct how the experimental results would look like if actin filaments were stable. While this approach shows that parts of the distribution of rupture forces of the actin–vimentin interaction is indeed masked by the breaking of actin, the result confirm that the quantitative parameters of the interaction does not change much between the different conditions.

In conclusion, our findings provide quantitative evidence that actin and vimentin filaments interact directly in the absence of crosslinking proteins, and these interactions persist across varying ionic conditions. By quantifying these interactions at the single-filament level, this study establishes a framework for linking molecular-scale mechanics to the emergent behavior of composite cytoskeletal networks. In this context, the direct coupling of actin and vimentin is likely to contribute to cellular mechanics and functions alongside protein-mediated crosslinking mechanisms.

## Author Contributions

S.Kö. conceived the project. S.Kö. and S.Kl. supervised the project; P.K. performed the experiments; P.K., S.L., K.B. and K.A.T.P. analyzed the data; S.L. developed the Bayesian approach; S.L., K.B. and K.A.T.P. performed Bayesian inference and simulations; P.K., S.Kö. and S.Kl. interpreted the results; P.K. and S.Kö. wrote a first draft of the manuscript, all authors contributed to writing the manuscript.

## Acknowledgments

We thank Charlotta Lorenz for training and consulting concerning the crossed-filaments experiments, Polina Malova, Susanne Bauch, and Kamila Sabagh for technical support. We thank them all for helpful discussions. This work was funded by the European Union’s Horizon Europe program under the Marie Skłodowska-Curie Actions (MSCA), Grant No. 101148781, and the Deutsche Forschungsgemeinschaft (DFG, German Research Foundation): Project-ID 449750155-RTG 2756, projects A3 and A7. This research was conducted within the Max Planck School Matter to Life, supported by the Dieter Schwarz Foundation and the German Federal Ministry of Research, Technology and Space (BMFTR) in collaboration with the Max Planck Society.

## Notes

### Competing Interest Statement

The authors have declared no competing interest.

