## Supplementary Material for "Interactions between single actin and vimentin filaments"

### Supplemental Methods

#### Crossed-filaments measurement geometry and force analysis

Figure S1 shows a schematic of the measurement geometry used in the optical tweezers experiments.

##### Interaction breaking forces

The interaction breaking forces are extracted following the procedure described in Refs. 1 and 2. Briefly, the total force acting at the interaction site (Figure S1a) is given by

$$F = F_{1y} + F_{2y}. \tag{1}$$

Only the force acting on bead 1 can be directly measured. Therefore, the total interaction force $F$ is calculated from the geometric configuration of the vertical filament relative to the horizontal filament. Because the forces in the $x$-direction cancel out ($F_{1x} = F_{2x}$), and defining the angles $\alpha_i$ (for $i = 1, 2$) of the deflected filament segments with respect to their original non-deflected orientations (Figure S1b), we have

$$\tan\alpha_1 = \frac{F_{1y}}{F_{1x}} = \frac{\Delta y}{l_1}, \quad \tan\alpha_2 = \frac{F_{2y}}{F_{2x}} = \frac{\Delta y}{l_2}. \tag{2}$$

From this relation, $F_{2y}$ can be expressed as

$$F_{2y} = F_{1y}\frac{l_1}{l_2}. \tag{3}$$

Using $l_1 + l_2 = l_{12}$ and Eq. (1), the interaction force $F$ becomes

$$F = F_{1y}\frac{l_{12}}{l_2}. \tag{4}$$

All measurements showing interactions are corrected by their respective geometric factors to obtain the true interaction force $F$ from the measured $F_{1y}$. The interaction breaking force ($F_b^{\mathrm{int}}$) is defined as the maximum corrected force observed just before the interaction breaks and the force drops to zero.

##### Actin breaking force of the vertical filament segments

The same equation applies when the vertical filament breaks.

##### Actin breaking force of the horizontal filament segments

**Left segment:** When the left actin segment ruptures (Figure S1c), the breaking force is given by

$$F_b^{\mathrm{actin}} = \sqrt{F_{1x}^2 + F_{1y}^2} \tag{5}$$

**Right segment:** When the right actin segment ruptures (Figure S1d), the breaking force can, in principle, be determined in the same way as for the left segment, but using the force on bead 2:

$$F_b^{\mathrm{actin}} = \sqrt{F_{2x}^2 + F_{2y}^2} \tag{6}$$

However, since the force experienced on bead 2 is not accessible, the breaking force cannot be extracted in this case. Therefore, this case is excluded from the data processing and quantitative analysis.

#### Two-bond model and Bayesian inference

##### Simulations of the two-bond model

The two-bond model describes the rupture of the actin–vimentin bond and of the actin filament as two competing stochastic processes that are subject to the same force $F(t)$. Simulating this model requires a force trajectory $F(t)$. We generate force trajectories in two ways: In the conceptual simulations of the model (Figure S7), this time-dependence of the force is obtained from an explicit polymer model for stretching of the filaments, while in the Bayesian inference and the posterior predictive simulations based on it (Figure 5 in the main text and Figures S8, S9), we directly use the experimental force trajectories.

The polymer model for the conceptual simulations is obtained, following Ref.;[2] briefly, in the experiments the vimentin filaments are pre-streched before the actin filament is pulled across the vimentin. We consider the initial increase in force observed after the bond is formed to be due to the entropic stretching of the actin filament. Thus, the end-to-end distance of the filament segment $x$ depends on the force $F$ via[3]

$$\frac{x}{d_{AC}} = \coth\left(\frac{2l_pF}{k_BT}\right) - \frac{k_BT}{2l_pF}, \tag{7}$$

where $d_{\mathrm{AC}}$ is the length of the actin segment between position of the bond on actin and bead 4 shown in Figure S1 and $l_p$ is the persistent length of actin. As bead 4 is moved with a constant velocity $v$, the end-to-end distance of the actin filament segment is $x = vt$. After a certain time $t_c$, the force is assumed to increase linearly over time. We obtain the force rate $w = dF/dt$ from the experimental force rate distribution. The transition time $t_c$ is obtained by matching the change in force $dF/dt$ due to the entropic stretching of the actin filament to the force rate $w$.[2] We consider the filament segment length $d_{\mathrm{AC}} = 4$ $\mu$m, which is roughly representative of the experimental filament segment lengths, however varies between different experiments and the force rate $w = 2$ pN/s which is the most probable force rate observed in the experiments. Persistence length of actin: $l_p = 10$ $\mu$m; the velocity of bead 4: $v = 0.3$ $\mu$m/s.

We emphasize again that in the Bayesian inference and the posterior predictive simulations described below, this polymer model of filament stretching is neither needed nor used and, thus, no assumptions about the stretching process are made. Instead we use directly the experimental time series $F(t)$, see below.

**Bayesian inference of bond parameters**

To obtain a complete kinetic description of the system that accounts for all experimental observations, we employ Bayesian inference within the two-bond model and infer the joint posterior over the four parameters

$$\theta = (r_{\mathrm{int}}, x_{\mathrm{int}}, r_{\mathrm{act}}, x_{\mathrm{act}})$$

from the full set of force–time trajectories obtained from both the crossed-filament experiment and the actin pulling experiment. In this framework, rupture is treated as a competing-risk process, in which actin failure censors the observation of actin–vimentin unbinding. By explicitly modeling both pathways, the inference incorporates this censoring and allows for simultaneous estimation of the kinetic parameters of both bonds.

The inferred model is furthermore used for posterior predictive simulations to remove the censoring effect. By suppressing actin rupture in the model, we predict the unbinding statistics of the actin–vimentin interaction in the absence of actin breaking and, thus, in the absence of masking.

Building on the Bayesian inference framework that we introduced recently[4] for single-bond systems, where bond parameters are inferred directly from full force–time trajectories, we extend it here to the system of two bonds in series with competing rupture pathways. The inference simultaneously targets the joint distribution of all kinetic parameters for both bonds: the interaction bond and the effective filament-breaking description.

The likelihood for the four parameters is constructed from the product of the two bonds' instantaneous unbinding rates. Note that this approach does not require a filament model to describe the time-dependent progression of forces. Instead, at each time point, the instantaneous load on the bonds is determined directly from experimental measurements, from which the hazard rates are derived. Labeling of the bond responsible for a breaking event is performed manually from confocal images, which clearly reveal whether the actin or the interaction bond breaks.

For a trajectory sampled at discrete time points $t_i$ with increments $\Delta t_i$, we construct the likelihood as a product over time bins. At each time step, the system can either remain bound, undergo actin rupture, or undergo interaction rupture. The instantaneous rates are evaluated at the measured force $F(t_i)$.

The probability of no rupture within a time step is

$$p_{\text{none}}(t_i) = \exp\big(-k_{\text{tot}}(t_i)\,\Delta t_i\big), \tag{8}$$

with $k_{\text{tot}}(t_i) = k_{\text{act}}(t_i) + k_{\text{int}}(t_i)$.

The probabilities for rupture events are

$$p_{\text{act}}(t_i) = 1 - \exp\big(-k_{\text{act}}(t_i)\,\Delta t_i\big), \tag{9}$$

$$p_{\text{int}}(t_i) = \big(1 - \exp(-k_{\text{int}}(t_i)\,\Delta t_i)\big)\exp\big(-k_{\text{act}}(t_i)\,\Delta t_i\big). \tag{10}$$

Using the observed label $y_i \in \{\text{none}, \text{act}, \text{int}\}$ for each time step, the log-likelihood is

$$\log \mathcal{L} = \sum_i \ell_i, \tag{11}$$

with per-step contributions

$$\ell_i = \begin{cases} -k_{\text{tot}}(t_i)\,\Delta t_i, & y_i = \text{none}, \\ \log\big(1 - \exp(-k_{\text{act}}(t_i)\,\Delta t_i)\big), & y_i = \text{act}, \\ \log\big(1 - \exp(-k_{\text{int}}(t_i)\,\Delta t_i)\big) - k_{\text{act}}(t_i)\,\Delta t_i, & y_i = \text{int}. \end{cases} \tag{12}$$

In addition, trajectories from single-actin pulling experiments are included, for which only the actin bond is present. In this case,

$$\ell_i^{\mathrm{actin}} = \begin{cases} -k_{\mathrm{act}}(t_i)\,\Delta t_i, & \text{no rupture,} \\ \log\big(1-\exp(-k_{\mathrm{act}}(t_i)\,\Delta t_i)\big), & \text{rupture.} \end{cases} \tag{13}$$

We assume uniform priors $P(\theta)$ over a broad positive parameter range. The posterior distribution is then given by

$$P(\theta \mid \mathrm{data}) \propto \mathcal{L}(\mathrm{data} \mid \theta)\, P(\theta), \tag{14}$$

and is sampled using an affine-invariant ensemble Markov Chain Monte Carlo method.[5]

**Posterior predictive simulations**

To assess the model and interpret the inferred parameters, we perform posterior predictive simulations. Parameter sets $\theta$ are sampled from the joint posterior, and rupture events are simulated along experimentally observed force–time trajectories using the time-dependent hazard $k_{\mathrm{tot}}(t)$. In cases where the predicted force trajectory survives longer than the experimental sample, we linearly extrapolate the force increase. At each time step $\Delta t$, rupture occurs with probability

$$1 - \exp\left(-k_{\mathrm{tot}}(t)\,\Delta t\right). \tag{15}$$

The corresponding breaking force is recorded at the first rupture event, yielding posterior predictive distributions of rupture forces that are compared to the experimental rupture force histograms, with which they are found to be consistent.

To isolate the intrinsic properties of the actin–vimentin bond, we perform counterfactual simulations in which actin rupture is suppressed by setting $r_{\mathrm{act}} = 0$, while keeping the interaction parameters unchanged. In this scenario, rupture can only occur via the actin–vimentin bond, allowing us to predict its unbinding statistics in the absence of masking by actin failure.

### Supplemental Figures

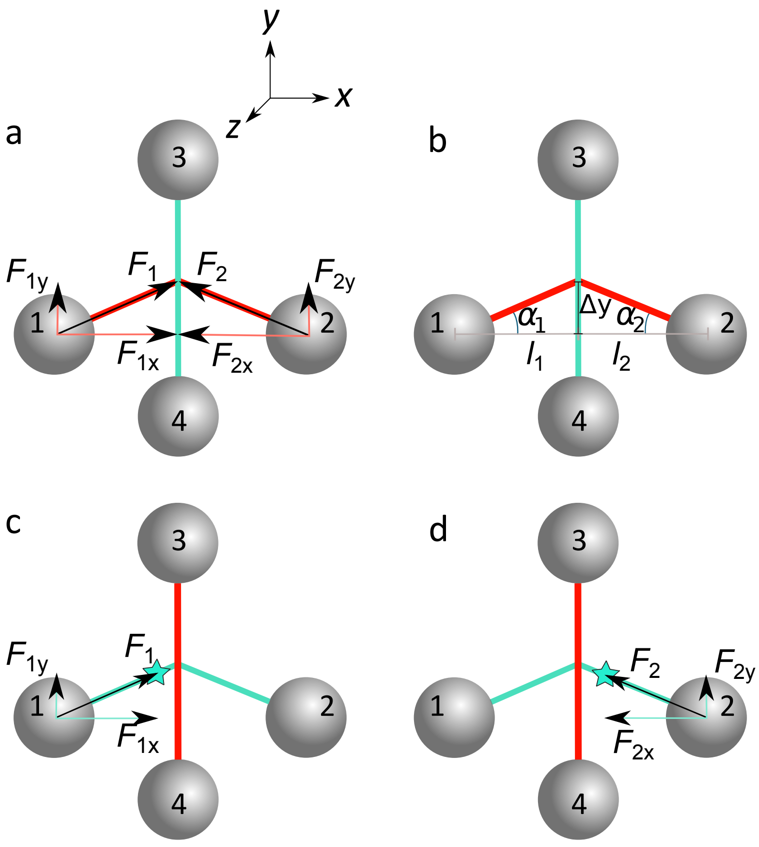


Figure S1: Schematic of the measurement geometry used to probe direct interactions between single vimentin and actin filaments. Actin is shown in cyan, vimentin in red. The "vertical" filament is moved along the $y$-axis within the $x$–$y$ plane to measure interactions ("upward" in the sketch). (a) Configuration 1; forces occurring when the filaments interact. The $x$-components of the forces cancel out. (b) Definition of geometric parameters (lengths and angles) used for data analysis. (c) Switched filament configuration (configuration 2) and forces occurring when the left actin segment ruptures. (d) Forces occurring in configuration 2 when the right actin segment ruptures. In this case the forces cannot be determined because the force detection on bead 2 is not possible.

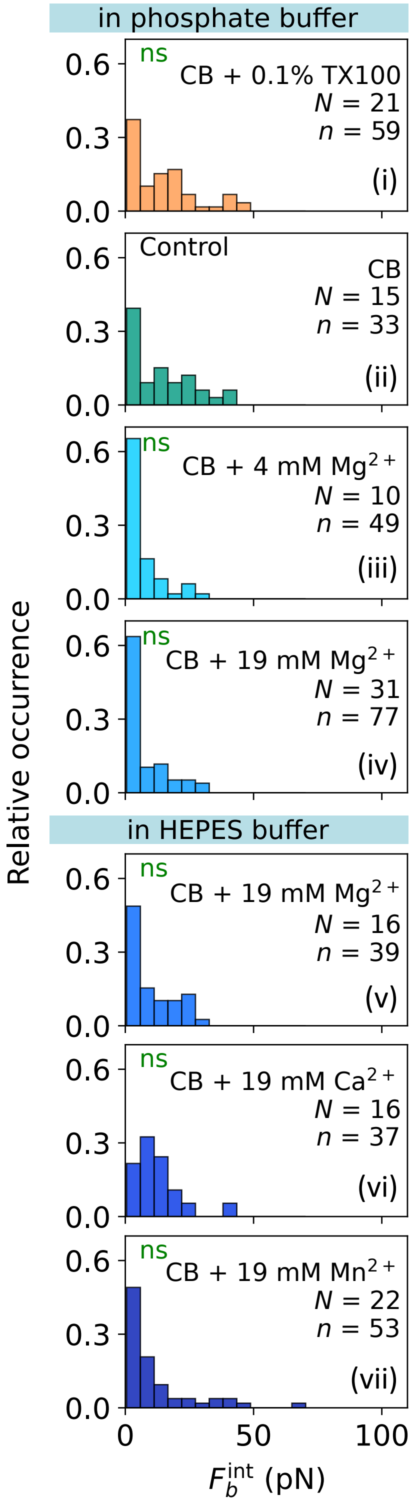


Figure S2: Histograms of interaction breaking forces at the indicated conditions (see information in the individual panels). Orange: addition of detergent; blue: addition of different types and concentrations of divalent ions. $N$ denotes the number of filament pairs, and $n$ denotes for the total number of interaction breaking events. To compare the distributions, $p$-values are calculated using a two-sample Kolmogorov–Smirnov (KS) test with a significance level of 5% followed by a Holm–Bonferroni correction. The statistical significance compared to CB is shown inside the panels; green indicates no significant (ns) difference.

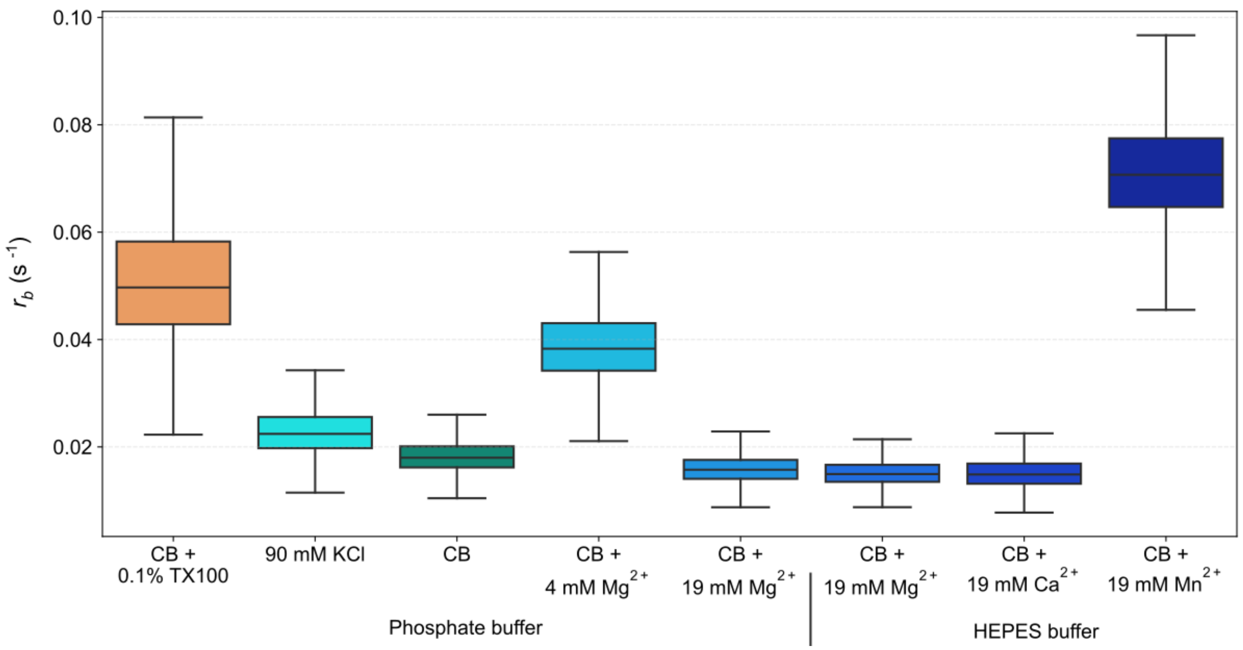


Figure S3: Actin–vimentin binding rates at the indicated conditions. Orange: addition of detergent; green: CB; blue: addition of different types and concentrations of divalent ions.

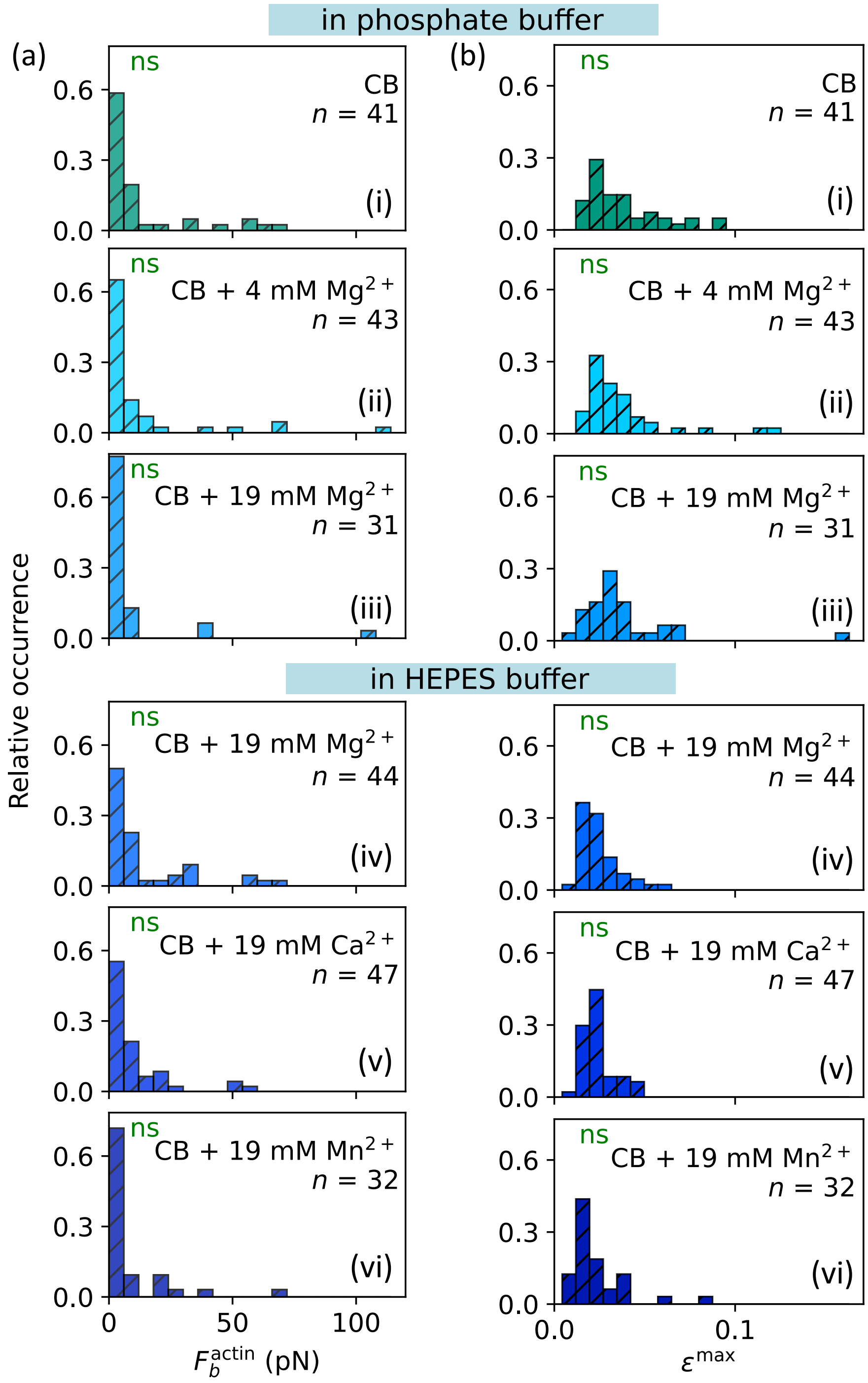


Figure S4: Histograms of (a) actin breaking forces and (b) maximum strains of single actin filaments prior to rupture. Data are obtained from single actin pulling experiments at the indicated conditions (see information in the individual panels). Green: CB; blue: addition of different types and concentrations of divalent ions. $n$ denotes the total number of actin breaking events. Differences between the breaking force distributions are assessed using the two-sample KS test with a significance level of 5%. The statistical significance compared to the measurements containing 90mM KCl shown in the main text is shown inside the panels. Differences in maximum strain between ionic conditions are assessed using the Kruskal–Wallis test followed by pairwise Mann–Whitney U tests with Holm–Bonferroni correction; green indicates no significant (ns) difference.

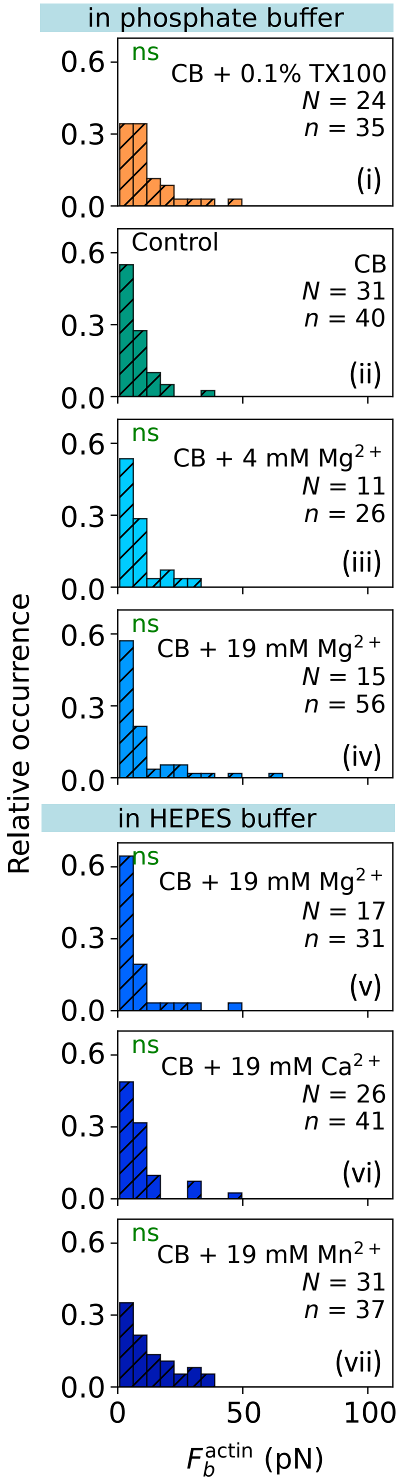


Figure S5: Histograms of actin breaking forces obtained from the crossed-filaments assay at the indicated conditions (see information in the individual panels). Orange: addition of detergent; green: CB; blue: addition of different types and concentrations of divalent ions. $N$ denotes the number of filament pairs, and $n$ denotes the total number of breaking events. To compare the distributions, $p$-values are calculated using the two-sample KS test with a significance level of 5% followed by a Holm–Bonferroni correction. The statistical significance compared to CB is shown inside the panels; green indicates no significant (ns) difference.

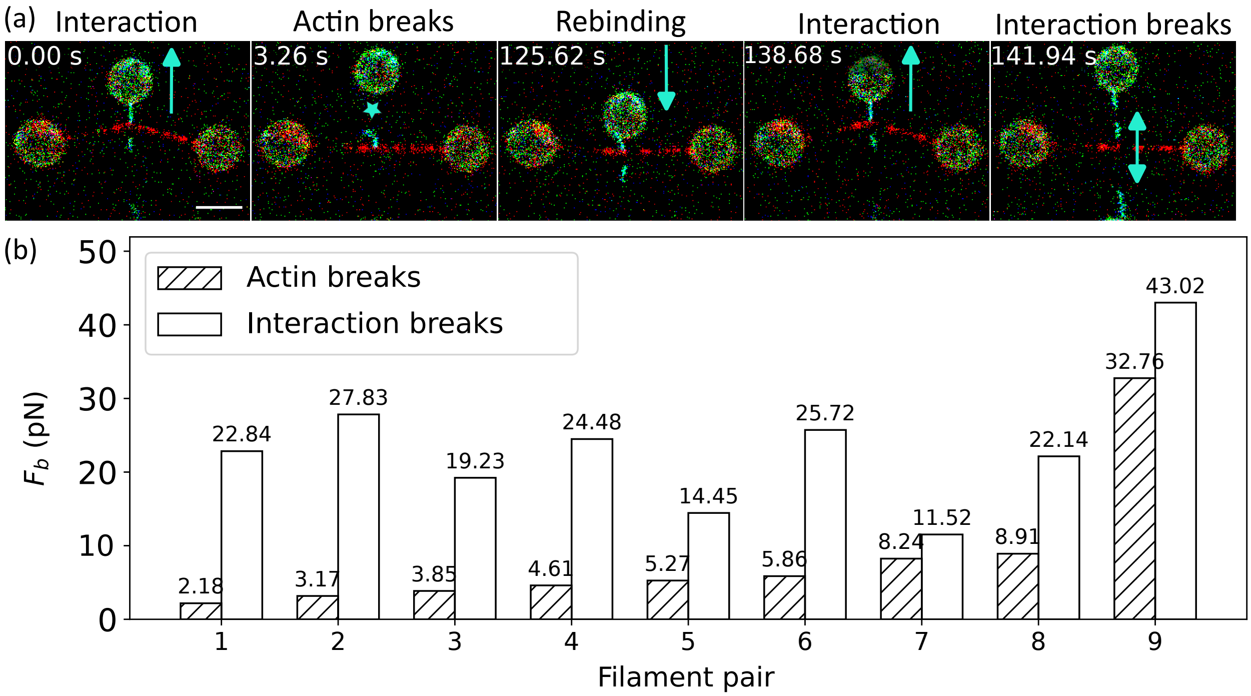


Figure S6: Actin breaking, rebinding and interaction breaking. (a) Time series of confocal fluorescence scans: when the vertical actin filament is displaced upwards along the $y$-axis, the filaments interact, causing the horizontal vimentin filament to bend. Upon further upward displacement, the actin filament breaks (cyan star). After subsequent downward displacement, the broken actin filament rebinds to the nearby bead. In the following upward displacement, the interaction breaks. Scale bar: 5 $\mu$m. (b) Bar plots comparing actin filament and interaction breaking forces across multiple filament pairs, demonstrating that after a filament broke, the subsequent interaction breaking force exceed the actin breaking force for the corresponding filaments pair.

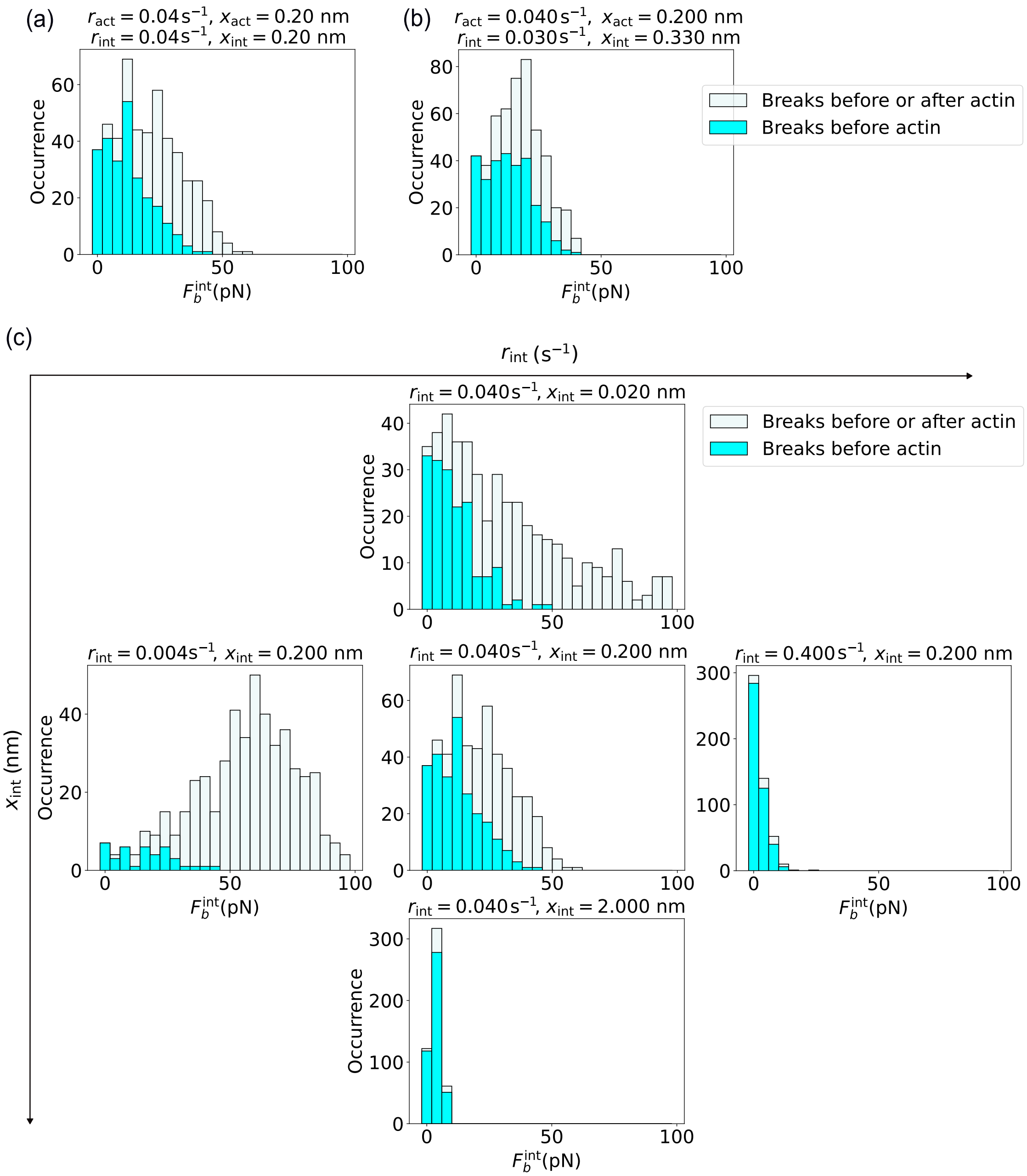


Figure S7: Masking of simulated interaction breaking force ($F_b$) distributions: Breaking forces are obtained from simulations of the two-bond model that follow the rupture of both bonds until the actin–vimentin interaction is broken, even if the actin filament breaks first, which is not possible in the experiments (500 simulated events per condition). As the bonds break independently, the full distribution of $F_b$ (light gray bars, including the cases where the interaction ruptures before the actin filament and after the actin filament) corresponds to the breaking force distribution in the case with stable actin. The subset where the interaction breaks first (cyan bars) gives the distribution observed in the quadruple optical tweezers experiment, where breaking of the actin filaments ends the experiment. (a) Scenario where the two bonds, i.e., the actin–vimentin interaction and the weak spot in actin, have the same parameters, (b) case with parameters near the maximum of the posterior distribution from Figure 5 in the main text. (c) Parameter dependence, varying the parameters of the interaction ($r_{\text{int}}, x_{\text{int}}$), while keeping the actin parameters fixed ($r_{\text{act}} = 0.04$ s$^{-1}$, $x_{\text{act}} = 0.2$ nm).

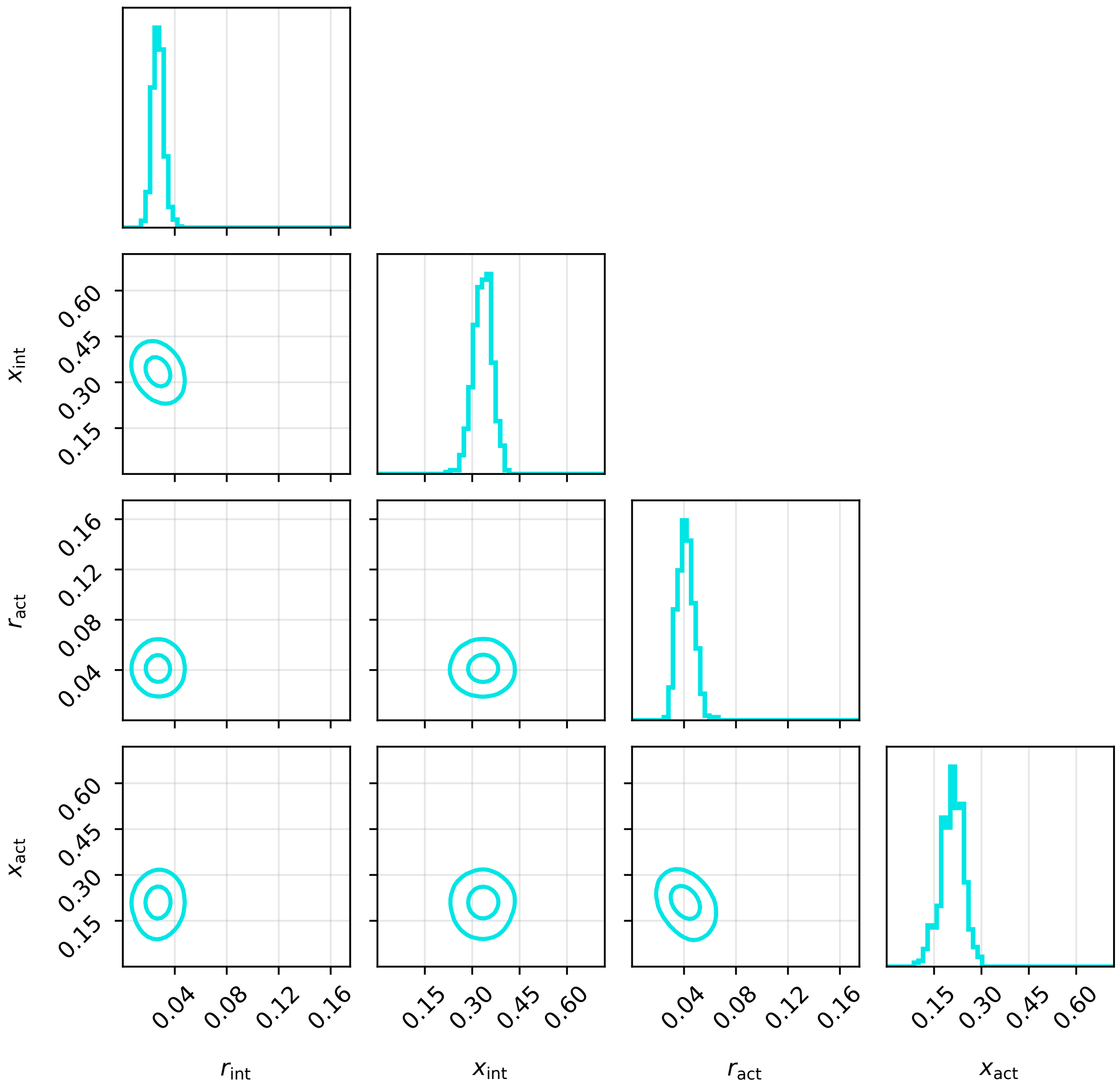


Figure S8: Corner plot of the full posterior distribution for the 90 mM KCl condition. The diagonals show the 1D marginal distributions of the 4 kinetic parameters. The off-diagonals are the pairwise 2D marginal distributions between all combinations of parameters. Drawn are the 95% and 50% highest density intervals. Marginal distributions are distinct from slices in that they integrate out the variables that are orthogonal to the displayed viewing direction, yielding a normalized probability distribution.

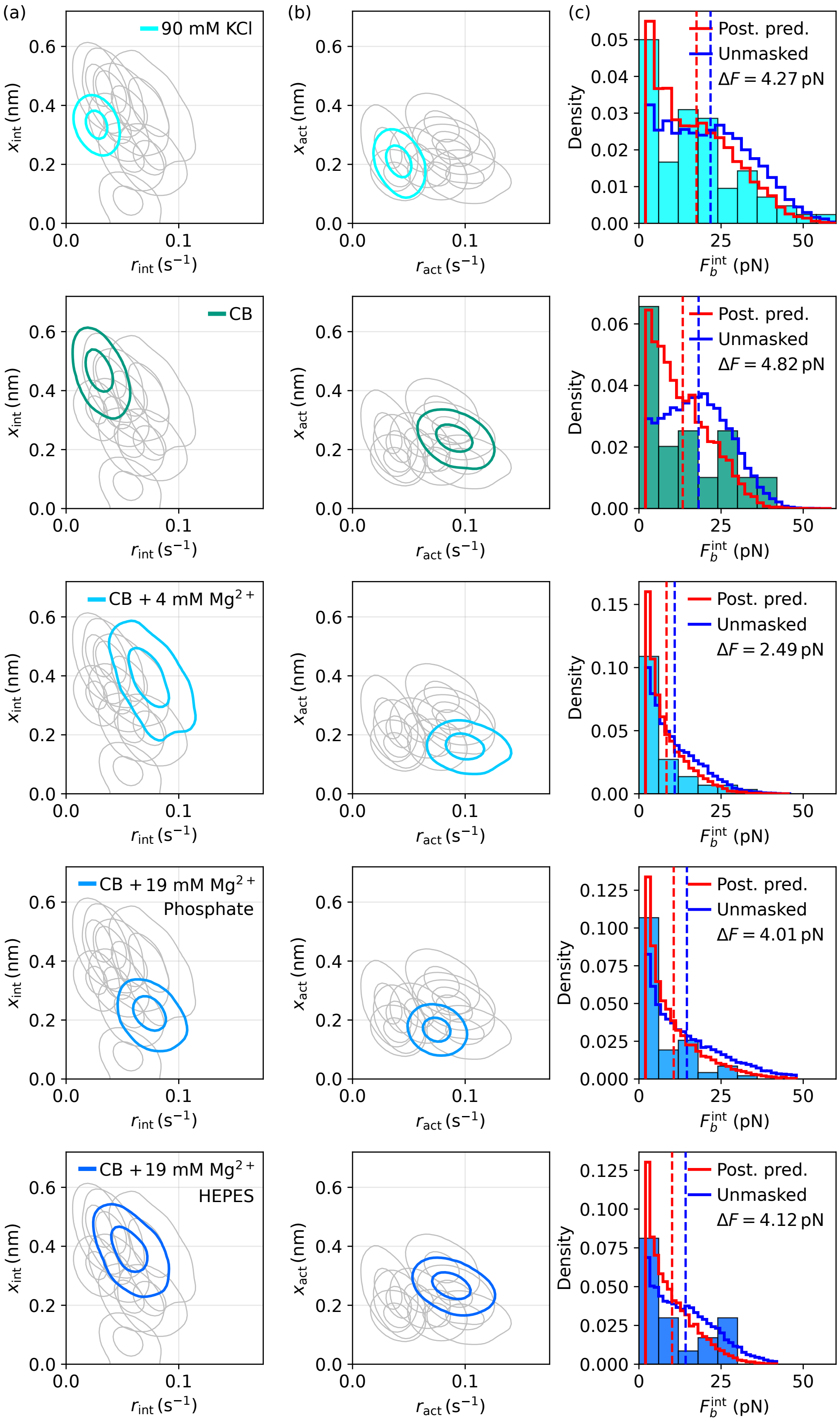

(a)
(b)
(c)
90 mM KCl
CB
CB + 4 mM Mg$^{2+}$
CB + 19 mM Mg$^{2+}$
Phosphate
CB + 19 mM Mg$^{2+}$
HEPES
$x_{\mathrm{int}}$ (nm)
$r_{\mathrm{int}}$ (s$^{-1}$)
$x_{\mathrm{act}}$ (nm)
$r_{\mathrm{act}}$ (s$^{-1}$)
Density
$F_b^{\mathrm{int}}$ (pN)
Post. pred.
Unmasked
$\Delta F = 4.27$ pN
$\Delta F = 4.82$ pN
$\Delta F = 2.49$ pN
$\Delta F = 4.01$ pN
$\Delta F = 4.12$ pN

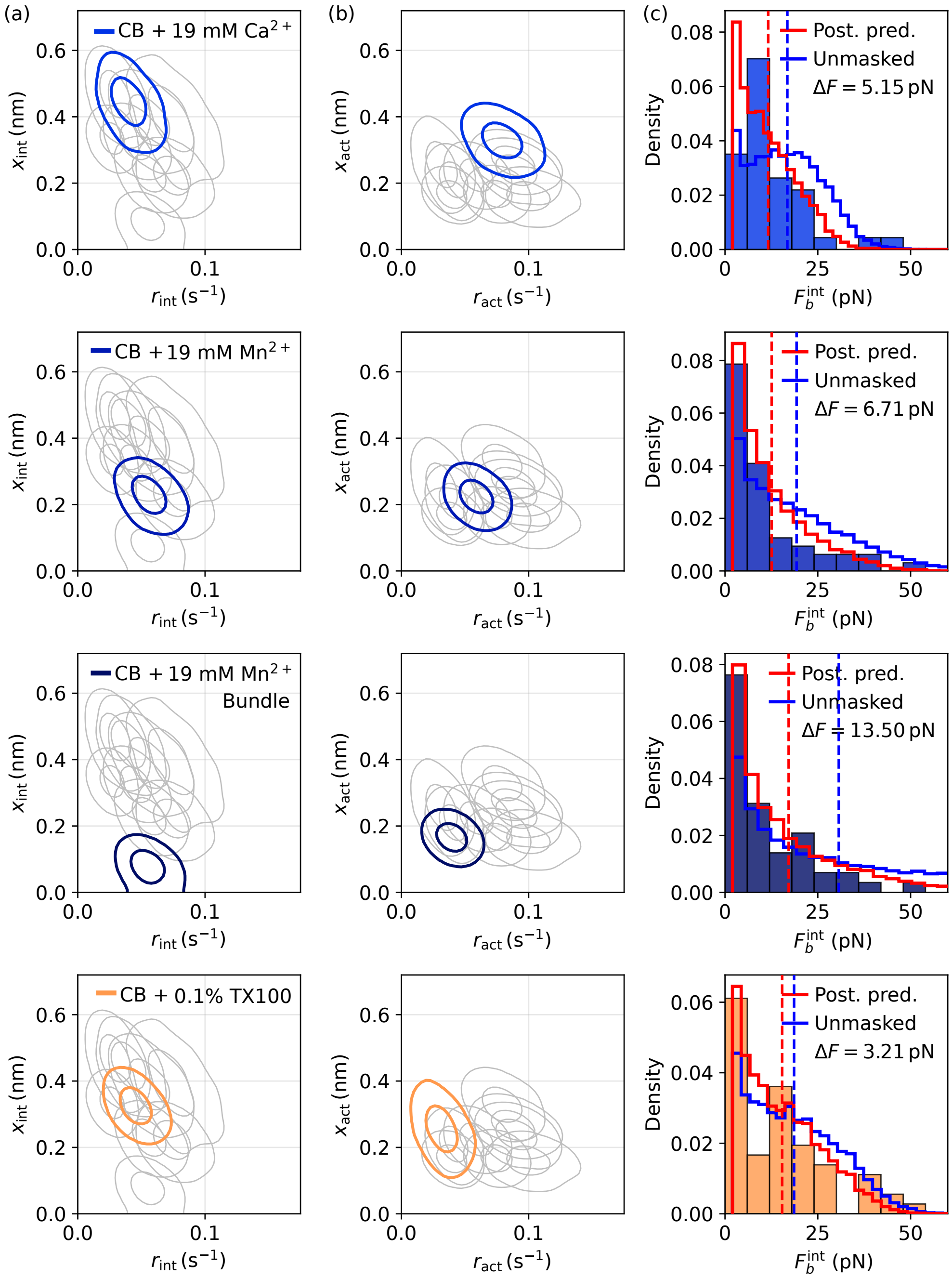


Figure S9: Bayesian inference results for all experimental datasets. Columns (a) and (b) show two slices of the 4D posterior distribution that describe all four kinetic bond parameters. Drawn are the 95% and 50% highest density intervals. Gray lines indicate the location of the posterior distributions of the other conditions. Each row corresponds to one buffer condition. Column (c) shows the empirical distribution of observed breaking forces as a histogram, along with the posterior predictive distribution of this observable from the inferred model (in red) and the counterfactual model with stabilized actin. The distributions show only the portion of results in which the actin does not break (which is not relevant to the stabilized actin prediction). The stabilized actin model shows a shift to higher breaking forces because actin breaking no longer masks long-lived interaction bonds. Vertical dashed lines indicate the prediction center of masses, separated by $\Delta F$. The 90 mM KCl posterior is the same as shown in the main text in Figure 5.

### Supplemental Movies

**Supplemental movie S1. Actin–vimentin interaction (i)** Time-lapse showing binding events between single actin and vimentin filaments in configuration 1 and subsequent interaction breaking.

**Supplemental movie S2. Actin–vimentin interaction (ii)** Time-lapse showing binding events between single actin and vimentin filaments in configuration 1 and subsequent actin breaking.

**Supplemental movie S3. Actin–vimentin interaction (iii)** Time-lapse showing binding events between single actin and vimentin filaments in configuration 2 followed up by interaction breaking and subsequent actin breaking.
